# Chemogenetic manipulation of microglia inhibits neuroinflammation and neuropathic pain in mice

**DOI:** 10.1101/2020.05.21.109538

**Authors:** Min-Hee Yi, Yong U. Liu, Kevin Liu, Tingjun Chen, Dale B. Bosco, Jiaying Zheng, Manling Xie, Lijun Zhou, Wenchun Qu, Long-Jun Wu

**Affiliations:** Department of Neurology, Mayo Clinic, Rochester, MN, 55905, USA; Robert Wood Johnson Medical School, Rutgers University, New Brunswick, NJ, 08854 USA; Department of Physiology and Pain Research Center, Zhongshan School of Medicine, Sun Yat-sen University, Guangzhou 510080, China; Department of Pain Medicine, Mayo Clinic, Jacksonville, FL, 32224, USA; Department of Neuroscience, Mayo Clinic, Jacksonville, FL, 32224, USA; Departments of Immunology, Mayo Clinic, Rochester, MN, 55905, USA

## Abstract

Microglia play an important role in the central sensitization and chronic pain. However, a direct connection between microglial function and the development of neuropathic pain *in vivo* remains incompletely understood. To address this issue, we applied chemogenetic approach by using CX_3_CR1^creER/+^:R26^LSL-hM4Di/+^ transgenic mice to enable expression of inhibitory Designer Receptors Exclusively Activated by Designer Drugs (Gi DREADD) exclusively in microglia. We found that microglial Gi DREADD activation inhibited spinal nerve transection (SNT)-induced microglial reactivity as well as chronic pain initiation and maintenance. Gi DREADD activation downregulated the transcription factor interferon regulatory factor 8 (IRF8) and its downstream target pro-inflammatory cytokine interleukin 1 beta (IL-1β). Using *in vivo* spinal cord recording, we found that activation of microglial Gi DREADD attenuated synaptic transmission following SNT. Our results demonstrate that microglial Gi DREADD reduces neuroinflammation, synaptic function and neuropathic pain after peripheral nerve injury. Thus, chemogenetic approaches provide a potential opportunity for interrogating microglial function and neuropathic pain treatment.

## Introduction

It has previously been demonstrated that neuroimmune interactions within the central nervous system (CNS) can mediate the pathogenesis of chronic pain (Ren and Dubner, 2010; Ji et al., 2016). In particular, microglia as CNS resident immune cells play a central role in the development of central sensitization and neuropathic pain (Ji and Suter, 2007; Inoue and Tsuda, 2018). Following peripheral nerve injury, microglia become activated and promote chronic pain by releasing a number of glial mediators that sensitize spinal neurons (Inoue et al., 2009; Zhuo et al., 2011). Indeed, specific ablation of microglia delayed the development of neuropathic pain (Peng et al., 2016). However, the direct connection between microglial function and neuropathic pain has not been clearly demonstrated *in vivo*.

G-protein-coupled receptors (GPCRs) and their downstream signaling regulate most physiological and pathological processes (Marinissen and Gutkind, 2001). To this end, Designer Receptors Exclusively Activated by Designer Drugs (DREADD) have been developed to precisely control various types of GPCR signaling (*i.e.*, Gq, Gs, Gi) using biologically inert agonists (Armbruster et al., 2007; Urban and Roth, 2015). DREADDs have been commonly used to modulate neuronal activity and interrogate neural circuitry of behaviors (Ilg et al., 2018). Recently, DREADD approaches in peripheral or central neurons were used to control pain transmission. For instance, activation of Gi DREADD in TRPV1^+^ DRG sensory neurons was able to decrease neuronal excitability, producing analgesia (Saloman et al., 2016), while activation of Gi DREADD in spinal dorsal horn GABAergic interneurons induced robust spontaneous nocifensive behaviors (Koga et al., 2017). Consequently, DREADD approaches offer a powerful tool for studying pain transmission and circuits.

While multiple studies have utilized neuronal DREADD approaches to modulate behaviors, few have investigated the effects of DREADD in glial cells. Previously, it has been shown that astrocytic Gq DREADD activation can elicit a number of physiological alterations, including changes in cardiovascular function, body temperature, activity-related behaviors, motor coordination (Agulhon et al., 2013; Sciolino et al., 2016) and improved cognitive performance (Adamsky et al., 2018). As for microglia, it has only recently been reported that AAV9-mediated DREADD expression in rat spinal microglia could affect morphine-induced nociceptive sensitivity (Grace et al., 2016) and peripheral nerve injury-induced neuropathic pain (Grace et al., 2018). However, there is no report on microglial DREADD approaches using genetic mouse models. Particularly, chemogenetic interrogations of mouse microglia and their downstream signaling pathways in chronic pain have so far been largely undetermined. Microglia express a number of GPCRs that are associated with Gi-signaling important for various microglia functions (Kettenmann et al., 2011). In particular, microglia signature P2Y12 receptor is a Gi-coupled GPCR. Our previous studies have found that P2Y12 receptors participate in microglia-neuron communication (Eyo and Wu, 2019) and in neuropathic pain (Gu et al., 2016a). However, it is still largely unknown how Gi signaling itself and its downstream pathways mediate microglial mechanism of pain.

In this study, we first used CX_3_CR1^creER/+^:R26^LSL-hM4Di/+^ transgenic mice to enable selective expression of Gi DREADD in microglia. Our results showed that chemogenetic inhibition of microglial function via Gi DREADD suppressed the development of neuropathic pain after spinal nerve transection (SNT). Moreover, we examined the molecular mechanisms by which microglial Gi DREADD reduced chronic pain. These results demonstrate that modulation of microglial Gi signaling directly impacts neuropathic pain pathogenesis and microglial chemogenetic approaches represent a potential opportunity for chronic pain treatment.

## Materials and methods

### Animals

7-12 weeks old male mice, unless otherwise indicated, were used in accordance with institutional guidelines as approved by the animal care and use committee at Mayo Clinic. C57BL/6J (000664) and R26^LSL-hM4Di^ (026219) (Zhu et al., 2016), and CX_3_CR1^CreER/CreER^ (021160) mice (Parkhurst et al., 2013) were purchased from Jackson Laboratory (Bar Harbor, ME). CX_3_CR1^CreER/CreER^ mice were crossed with R26^LSL-hM4Di^ mice to obtain CX_3_CR1^creER/+^:R26^LSL-hM4Di/+^ mice. These mice were assigned to an experimental group randomly within a litter. Experimenters were blind to drug treatments.

### Gi DREADD activation

To induce Gi DREADD expression in CX_3_CR1^+^ cells, 150 mg/kg tamoxifen (TM; T5648, Millipore-Sigma, Burlington, MA) in corn oil (20 mg/mL) was administered via *i.p.* injection twice daily for 2 days. To activate Gi DREADD, 5 mg/kg Clozapine N-oxide (CNO; 16882, Cayman Chemical, Ann Arbor, MI) was administered *i.p.* daily for 3 days. WT (C57BL/6J) mice receiving TM and CNO and CX_3_CR1^creER/+^:R26^LSL-hM4Di/+^ receiving TM and vehicle (but not CNO) were used for controls.

### Spinal nerve transection (SNT) induced pain induction

SNT surgery was performed under 2% isoflurane anesthesia. Briefly, an incision was made along the mid-line of the lumbar spine. The left paraspinal muscles in front of the pelvic bone were separated to expose the L5 transverse process. The L5 transverse process was then removed to expose the L4 spinal nerve. The L4 spinal nerve was separated, transected and removed 1-1.5 mm from the end to DRG. The wound was then irrigated with sterile PBS and sutured.

### Behavioral assessments

Mechanical allodynia was assessed by measuring paw withdrawal threshold test via von Frey filaments (0.04-2 g). Briefly, mice were placed on an elevated metal grid and filaments of increasing weight were applied to the plantar surface at a vertical angle for up to 3 s. Fifty percent withdrawal threshold values were determined using the up-down method (Chaplan et al., 1994).

Thermal hyperalgesia was assessed by measuring paw withdrawal latency to radiant heat stimuli. Briefly, mice were placed in elevated chambers with a plexiglass floor and allowed to habituate for 20 min. A radiant heat source was then applied to the center of the plantar surface of the hind paw four times with at least 3-min intervals. The average withdrawal latency of the four trials was recorded as the response latency.

### Tissue collection

At various time points post SNT, mice were deeply anesthetized and perfused transcardially with 20 mL PBS followed by 20 mL of cold 4% paraformaldehyde (PFA) in PBS. Spinal cords were then removed and post-fixed with 4% PFA for 6 h at 4°C. The samples were then transferred to 30% sucrose in PBS overnight. 15 μm thick sections were prepared via cryosection and applied to charged glass slides. Slides were then stored at -20°C until use.

### Fluorescent immunostaining

Tissue slides were blocked with 5% goat serum in 0.3% triton X-100 (Sigma) in PBS buffer for 60 min, and then incubated overnight at 4°C with primary antibody for rat anti-HA (1:200; 12013819001, RRID:AB_390917, Roche, Basel, Switzerland), rabbit anti-GFP (1:200; ab290, RRID:AB_303395, Abcam, Cambridge, United Kingdom), rabbit anti-Iba1 (1:500; 019-19741, RRID:AB_839504, FUJIFILM Wako Chemicals USA, Richmond, VA), rat anti-CD11b (1:200; 101202, RRID:AB_312785, BioLegend, San Diego, CA), mouse anti-IRF8 (1:400; sc-365042, RRID:AB_10850401, Santa Cruz Biotechnology, Dallas, Texas) and mouse anti-IL-1β (1:400; 12242, RRID:AB_2715503, Cell Signaling Technology, Danvers, MA). The sections were then incubated for 60 min at room temperature, with goat anti-rat (1:500; A-11006, RRID:AB_2534074, Thermo Fisher Scientific, Waltham, MA), goat anti-rabbit (1:500; A-11035, RRID:AB_2534093, Thermo Fisher Scientific) or goat anti-mouse secondary antibodies (1:500; A-11029, RRID:AB_2534088, Thermo Fisher Scientific). The sections were then mounted with Fluoromount-G (Southern Biotech) and fluorescent images were obtained with an EVOS FL Imaging System (Thermo Fisher Scientific).

### Western blot analysis

Lumbar 4-5 spinal dorsal horns were collected at various time points and protein was extracted. 50 µg of protein from each group was then loaded and separated by SDS-PADGE, transferred to a PVDF membrane, blocked with 5 % skim milk in TBST, and incubated overnight with primary antibodies at 4°C. Primary antibodies include, mouse anti-IRF8 (1:1000), mouse anti-IL-1β (1:1000), and GAPDH (1:1000; sc-32233, RRID:AB_627679, Santa Cruz Biotechnology). Membranes were incubated with horseradish peroxidase-conjugated goat anti-mouse IgG (1:3,000; 115-035-003, RRID:AB_10015289, Jackson ImmunoResearch Labs, West Grove, Pennsylvania) for 1 hr at room temperature. Membranes were then treated with West Pico substrate (34078, Thermo Fisher Scientific) and chemiluminescence signal was detected with a G:BOX Chemi XRQ gel doc (Syngene, Frederick, MD). Optical density of each band was then determined using Fiji, (NIH).

### Sholl analysis

Z-stack confocal live microglia images (30 μm) were acquired at 3-μm intervals using a 40× objective of confocal microscope (LSM510, Zeiss). Consecutive Z-stack Images were converted to a maximum intensity projection image using Fiji software. Using the Image5D plugin (Fiji, NIH), z-stack images were condensed into a maximum intensity projection image over which concentric circles were drawn (concentric circles plugin, fiji), centered on the soma, beginning at 0.5 μm radii and increasing 0.1 μm with every circle. Sholl analysis was manually performed for each cell by counting the number of intersections between microglia branches and each increasing circle to create a Sholl plot. Additional measures to characterize each cell included the number of branch number, length (manual count) and cell soma area (Fiji Analysis).

### *In vivo* electrophysiology recording in the spinal cord dorsal horn

Under anesthesia, a T12-L1 laminectomy was performed to expose the lumbar enlargement of the spinal cord and the dura was removed. The left sciatic nerve was then exposed for electrical stimulation with a bipolar platinum hook electrode. A test stimulus (0.5 ms duration, every 1 min, at C-fiber intensity) was then delivered to the sciatic nerve. C-fiber evoked fEPSP was recorded from the dorsal horn with a glass microelectrode filled with 0.5 M sodium acetate (impedance 0.5–1 MU). An A/D converter card (M-Series PCI-6221, National Instruments, Austin, TX) was used to digitize and store data at a sampling rate of 10 kHz. C-fiber evoked fEPSP was determined with WinLTP Standard Version (WinLTP Ltd., Bristol, United Kingdom). For each experiment the average amplitude of five consecutive fEPSP was recorded at 30 sec intervals. The mean amplitude of the responses before CNO administration served as baseline. To observe the effect of DREADD activation on C-fiber evoked field potentials, CNO was injected *i.p.* 30 mins after stable recording of C-fiber evoked field potentials.

### Statistical analysis

Pain behaviors were analyzed using two-way ANOVA to test for main effects between groups followed by post-hoc testing for significant differences by day. Two group analyses utilized Student’s t-test. Data were represented as mean ± SEM. All statistical analyses were performed using Prism6 (<u>GraphPad</u>, San Diego, CA). Level of significance is indicated with *p < 0.05, ** p < 0.01, *** p < 0.001, **** p < 0.0001

## Results

### Cre-dependent expression of hM4Di Gi DREADD in CX_3_CR1^+^ cells

The chemokine receptor CX_3_CR1 is highly expressed by microglia in the CNS and cells of mononuclear origin in the periphery (Jung et al., 2000). To study the role of CX_3_CR1^+^ cells in neuropathic pain, we first generated CX_3_CR1^creER/+^:R26^LSL-hM4Di/+^ mice to enable tamoxifen (TM) inducible Cre-mediated expression of Gi DREADD in CX_3_CR1^+^ cells (**Fig. 1A-B**). To confirm Gi DREADD expression, we stained spinal cord sections for mCitrine and HA tag which are linked to hM4Di expression (Zhu et al., 2016). We found that mCitrine and HA tag expression was induced by TM injection (**Fig. 1C-D**). To validate that Gi DREADD expression was specific to CX_3_CR1^+^ microglia, spinal and brain slices were examined for co-localization of Iba1 with HA tag in mice with or without TM treatment (**Fig. 1C**). As expected, in mice receiving TM, there was near total co-localization of HA with Iba1 in spinal dorsal horn (SDH) but not in control group without TM treatment (**Fig. 1C**). Similarly, mCitrine and HA co-localization was observed in cortex and hippocampus of TM treated group (**Fig. 1D**). In addition, Iba1 positive cell co-localized with HA in the cortex of with TM mice unlike the without TM mice. (**Fig. 1D**). These results indicate that we were able to induce Cre-dependent expression of Gi DREADD specifically in CX_3_CR1^+^ cells in the CNS.

**Fig. 1.**
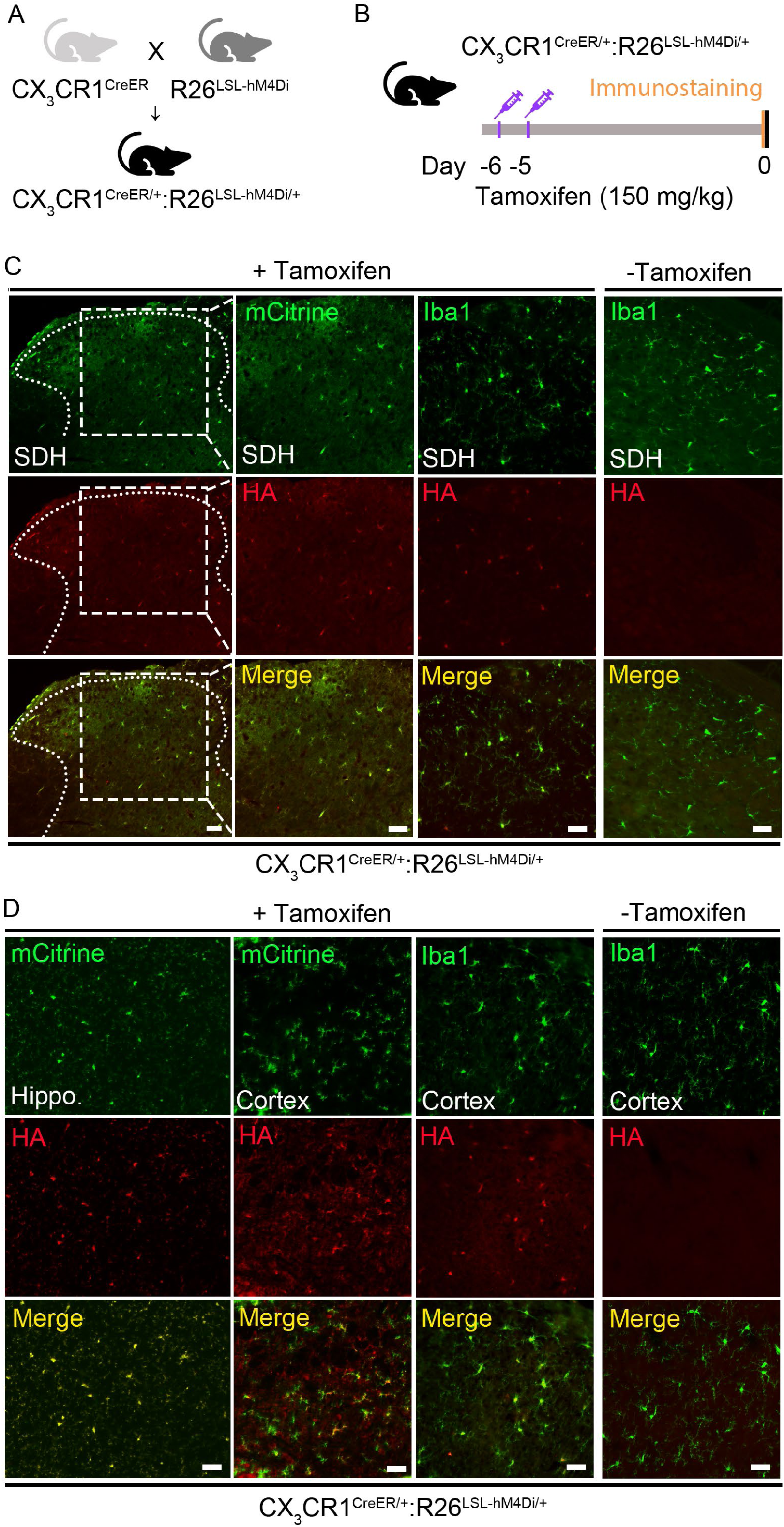
CX_3_CR1^creER/+^:R26^LSL-hM4Di/+^ transgenic mice enable selective expression of Gi DREADD in microglia. (A) Generation of CX_3_CR1^creER/+^:R26^LSL-hM4Di/+^ transgenic mice. (B) Timeline of experimental procedures for tamoxifen (TM) injection and immunostaining. (C) Representative images of mCitrine (green) and HA immunostaining (red) in spinal cord dorsal horn (SDH) of CX_3_CR1^creER/+^:R26^LSL-hM4Di/+^ mice after TM. Co-labeling Iba1 (green) and HA (red) showed that HA expression was co-localized with Iba1^+^ cells in SDH region of CX_3_CR1^creER/+^:R26^LSL-hM4Di/+^ mice with TM, but not in CX_3_CR1^creER/+^:R26^LSL-hM4Di/+^ mice without TM. Scale bar, 40 μm. n=5 mice/group. (D) Immunofluorescence images of the hippocampus and cortex labeled with mCitrine and HA in CX_3_CR1^creER/+^:R26^LSL-hM4Di/+^ mice after TM injection show expression of Gi DREADD. Co-staining of HA (red) with Iba1 (green) in brain cortex of CX_3_CR1^creER/+^:R26^LSL-hM4Di/+^ mice with TM indicates co-localization which is absent without TM injection. Scale bar, 40 μm. n=5 mice/group.

### Activation of Gi DREADD in CX_3_CR1^+^ cells attenuates chronic pain after SNT

To address the function of Gi DREADD in CX_3_CR1^+^ cells in chronic pain, we used SNT model to induce neuropathic pain in three experimental groups: WT mice with DREAD ligand CNO (WT+CNO); CX_3_CR1^creER/+^:R26^LSL-hM4Di/+^ mice with vehicle (Gi DREADD+Vehicle), and CX_3_CR1^creER/+^:R26^LSL-hM4Di/+^ mice with CNO (Gi DREADD+CNO). We injected TM into these mice starting 6d before SNT to induce cre-medicated Gi DREADD expression in CX_3_CR1^+^ cells, then either CNO or vehicle was administered starting 3d before SNT (pre-SNT) (**Fig. 2A**). We compared SNT induced chronic pain behaviors in mice with or without Gi DREADD activation. Our results showed that pre-SNT Gi DREADD activation significantly attenuated mechanical allodynia for a 4 days period in the ipsilateral side of SNT in Gi DREADD+CNO group, when compared to Gi DREADD+Vehicle control mice. CNO itself has no effect on chronic pain induced by SNT in WT mice (**Fig. 2B**). In addition, no difference was observed between any groups in contralateral response (**Fig. 2C**). These results indicate that pre-SNT activation of Gi DREADD in CX_3_CR1^+^ cells can delay the development of neuropathic pain.

**Fig. 2.**
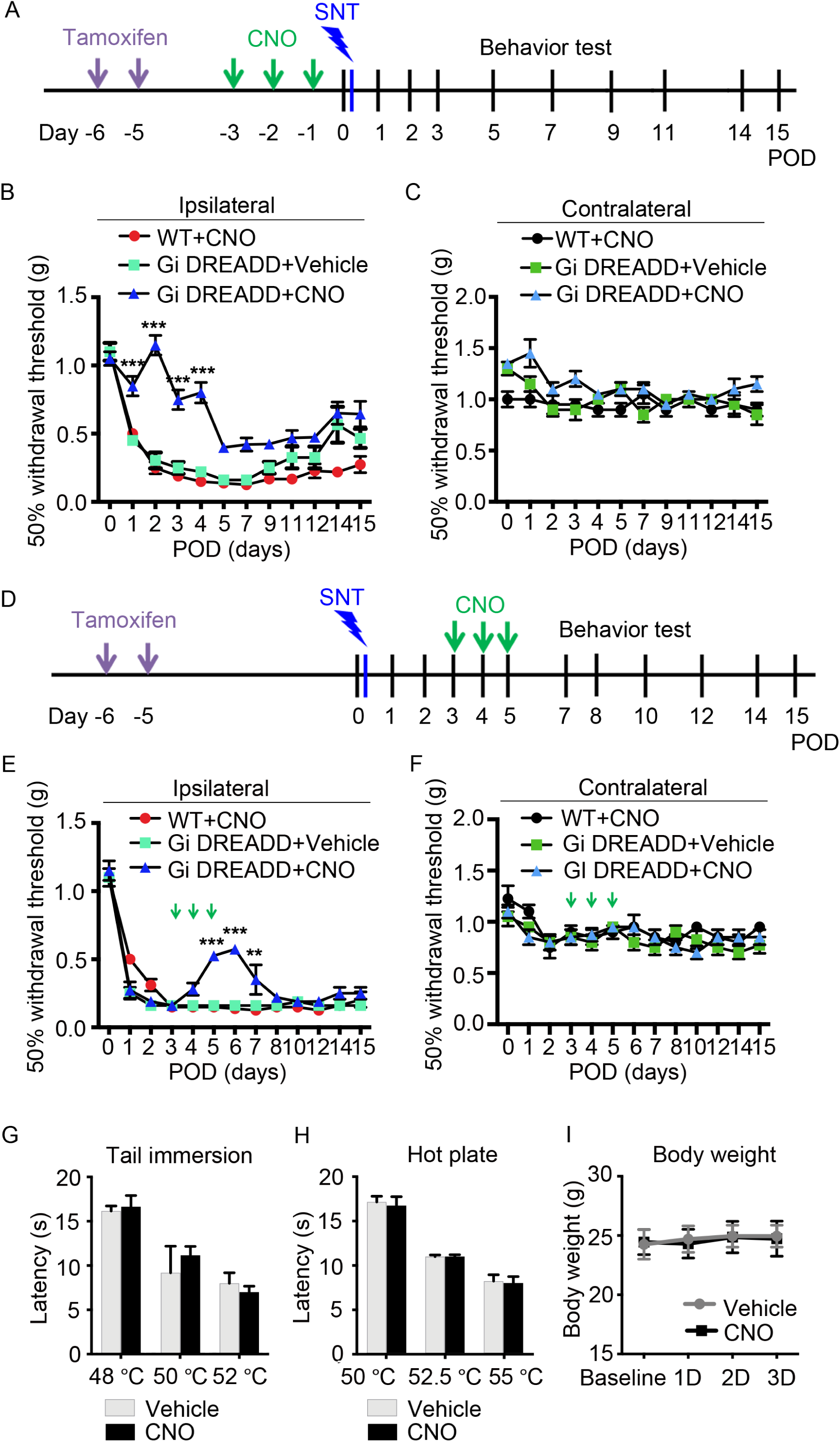
Activation of Gi DREADD in CX_3_CR1^+^ cells attenuates chronic pain after SNT. (A) Timeline of experimental procedures for TM injection, pre-SNT CNO administration, SNT surgery, and pain behavioral test. (B, C) Measurement of mechanical hypersensitivity in WT mice with CNO (WT+CNO), CX_3_CR1^creER/+^:R26^LSL-hM4Di/+^ mice with vehicle (Gi DREADD+Vehicle), and CX_3_CR1^creER/+^:R26^LSL-hM4Di/+^ mice with CNO (Gi DREADD+CNO). Activation of Gi DREADD in CX_3_CR1^+^ cells following SNT showed that initiation of mechanical hypersensitivity was delayed when compared to control groups ipsilaterally (B), but no significant change was observed contralaterally (C). Data represented as mean ± SEM, n=5-8 mice/group, *** p < 0.001. (D) Timeline of experimental procedures for TM injection, SNT surgery, post-SNT CNO administration, and pain behavioral test. (E) Mechanical hypersensitivity showing that post-SNT activation of Gi DREADD significantly reversed the pain behaviors ipsilaterally in Gi DREADD+CNO group when compared to control groups. (F) However, no difference was found between the three groups in the contralateral side. Data represented as mean ± SEM, n=8 mice/group, ** p < 0.01, *** p < 0.001. (G, H) Effect of Gi DREADD on acute pain was tested by analyzing tail withdrawal latencies in tail immersion test (G) and hot plate test (H). (I) Body weight was measured in CX_3_CR1^creER/+^:R26^LSL-hM4Di/+^ mice with vehicle (grey) or with CNO treatment (black). The results show no significant change between the vehicle and CNO treated groups. Data represented as mean ± SEM, n=5 mice/group.

We also wanted to understand the effect of Gi DREADD activation in CX_3_CR1^+^ cells on the maintenance of neuropathic pain. Therefore, we administered CNO at postoperative days (POD) 3 to POD5 after SNT (post-SNT) and examined chronic pain behaviors (**Fig. 2D**). We found that post-SNT Gi DREADD activation significantly attenuated mechanical allodynia starting POD5 in Gi DREADD+CNO group, but not in Gi DREADD+Vehicle control or WT+CNO groups (**Fig. 2E**). Interestingly, the effect of post-SNT Gi DREADD activation only lasted for 3 days (**Fig. 2E**). No significant difference between all three groups was observed in contralateral side of SNT (**Fig. 2F**). Therefore, post-SNT activation of Gi DREADD in CX_3_CR1^+^ cells temporally reversed the maintenance of mechanical allodynia induced by peripheral nerve injury.

To ensure that these observations were specific to SNT-induced chronic pain but not acute pain, we examined the effects of Gi DREADD activation by CNO on tail immersion (**Fig. 2G**) and hot plate test in Gi DREADD+CNO groups and Gi DREADD+Vehicle groups (**Fig. 2H**). For both assays no difference between two groups was observed. Gi DREADD activation also had no apparent effect on body weight (**Fig. 2I**). Taken together, our results indicate that both pre- and post-SNT activation of Gi DREADD in CX_3_CR1^+^ cells can suppress the development and the maintenance of neuropathic pain.

### Activation of Gi DREADD specific to microglia inhibits neuropathic pain

Previous studies have demonstrated that CX_3_CR1^+^ cells, including microglia, monocytes, NK cells are all critically involved in neuropathic pain (Ji et al., 2016). To exclusively evaluate the function of microglial Gi DREADD in chronic pain, we injected TM into CX_3_CR1^creER/+^:R26^LSL-hM4Di/+^ mice 28d before SNT (**Fig. 3A**). This is to take advantage of the significant difference in turnover rates between microglia and peripheral mononuclear cells, allowing for exclusive expression of Gi DREADD within microglia (Parkhurst et al., 2013). We found pre-SNT activation of Gi DREADD by CNO significantly reduced mechanical allodynia following SNT in ipsilateral hindpaws of Gi DREADD+CNO group, but not in WT+CNO mice or in Gi DREADD+Vehicle group (**Fig 3B**). No significant difference between all three groups was observed in the contralateral side **(Fig 3C**). Thus, pre-SNT activation of microglial Gi DREADD delays the development of mechanical allodynia following SNT.

**Fig. 3.**
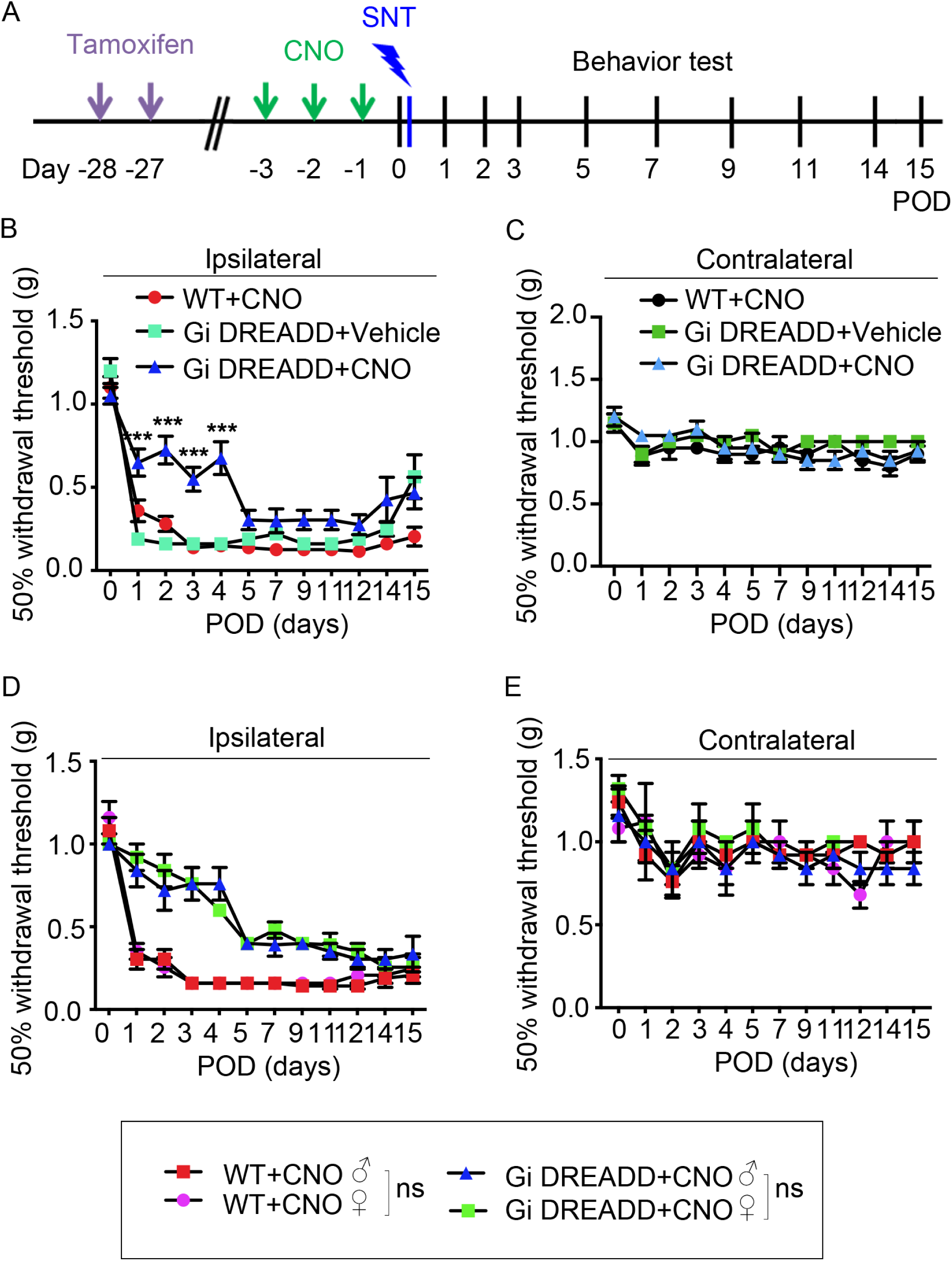
Activation of Gi DREADD in microglia delays the development of chronic pain after SNT in both male and female mice. (A) Timeline of experimental procedures. (B, C) Pre-SNT activation of microglial Gi DREADD delayed mechanical hypersensitivity in Gi DREADD+CNO group when compare to control groups (WT+CNO or Gi DREADD+Vehicle) ipsilaterally (B), but no significant change was observed contralaterally (C). Data represented as mean ± SEM, n=8 mice/group, *** p < 0.001. (D) Pain behavioral tests in the ipsilateral side showed pre-SNT activation of microglial Gi DREADD reduced mechanical hypersensitivity in both male and female CX_3_CR1^creER/+^:R26^LSL-hM4Di/+^ mice but not WT mice. There is no significant difference in pain response between male and female mice. (E) No difference was observed between all groups contralaterally. Data represented as mean ± SEM, n=5 mice/group.

It has been reported that microglia play more important role in pathological pain in male than in female mice (Sorge et al., 2015; Chen et al., 2018). Therefore, we wanted to examine whether microglial Gi DREADD function in neuropathic pain was also sex dependent. To this end, we compared the CNO effect on pain behavior in male and female CX_3_CR1^creER/+^:R26^LSL-hM4Di/+^ mice (or WT mice as control) following SNT. Interestingly, we found no difference in the inhibitory effect of microglial Gi DREADD on pain response between male and female mice (**Fig 3D**), and also no significant difference in contralateral side (**Fig. 3E**). Together, our results highlight that microglia specific Gi DREADD activation delays the development of neuropathic pain in both male and female mice following SNT.

### Activation of Gi DREADD suppresses SNT-induced microglial activation

There is significant proliferation and activation of spinal microglia after peripheral nerve injury (Zhang et al., 2008; Inoue and Tsuda, 2018). As such, we first examined microglial numbers at POD3 after SNT (**Fig. 4A**), when it is the peak of spinal microglia proliferation in response to peripheral nerve injury (Gu et al., 2016b; Tashima et al., 2016). Consistently, the number of microglia labeled by Iba1 immunostaining in the ipsilateral dorsal horn was dramatically increased following SNT. Further, we found that pre-SNT Gi DREADD activation by CNO in CX_3_CR1^creER/+^:R26^LSL-hM4Di/+^ mice largely reduced SNT-induced microglial numbers compared with the vehicle group (**Fig. 4B, C**). Using CD11b immunostaining, we observed the similar inhibition of microglial numbers by pre-SNT CNO treatment (Data not shown). Thus, pre-SNT activation of microglial Gi DREADD signaling inhibits the increase of spinal microglial numbers induced by peripheral nerve injury.

**Fig. 4.**
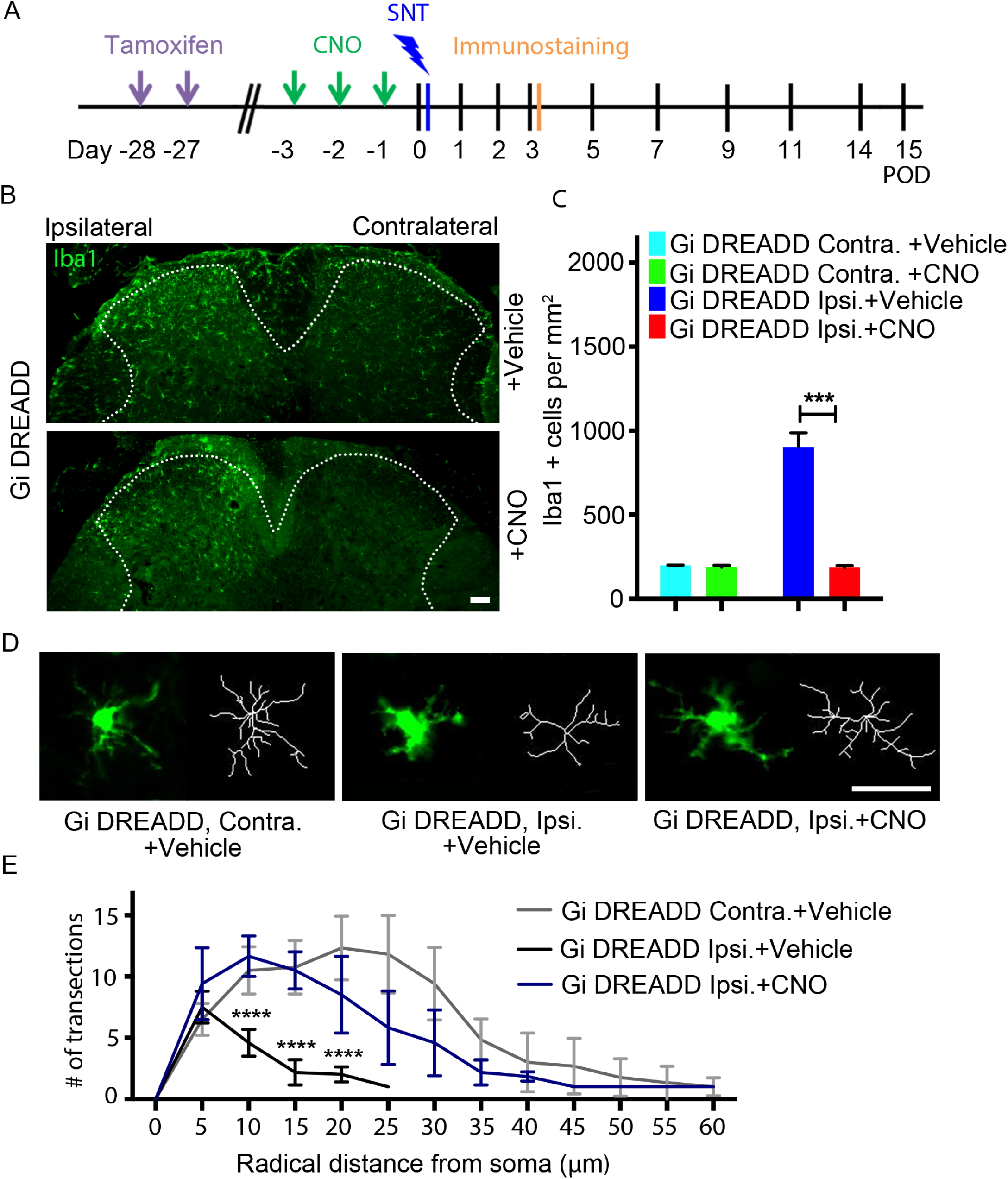
Microglial Gi DREADD activation reduces microglia activation. (A) Timeline of experimental procedures. (B) Representative immunostaining images show that the number of Iba1^+^ microglia was reduced in the ipsilateral dorsal horn at POD3 by pre-SNT activation of microglial Gi DREADD in CX_3_CR1^creER/+^:R26^LSL-hM4Di/+^ mice. Scale bar, 40 μm. Data represented as mean ± SEM, n=8 mice/group. (C) Summarized data showing microglial numbers were reduced by CNO treatment but not in vehicle group. Scale bar, 40 μm. Data represented as mean ± SEM, n=8 mice/group, *** p < 0.001. (D) Representative single microglia images in the spinal cord following SNT by Iba1 immunostaining and after being skeletonized. (E) Sholl analysis showing CNO treatment increased the complexity of microglia compared with vehicle group. Data represented as mean ± SEM, n=5 mice/group, **** p < 0.0001.

It has been shown that microglial morphology has been directly related to their activation state. For instance, microglial processes are visibly shorter and less complex within a day of injury (Inoue and Tsuda, 2009; Gu et al., 2016a). Therefore, here we examined the effects of Gi DREADD activation on microglial morphology in CX_3_CR1^creER/+^:R26^LSL-hM4Di/+^ mice following SNT. Using Sholl analysis, we compared the complexity of spinal microglia after CNO or vehicle treatment (**Fig. 4D**). Indeed, we found that pre-SNT Gi DREADD activation reversed SNT-induced morphological change (**Fig. 4D, E**). Consequently, these results further indicate that Gi DREADD inhibits microglial activation in response to SNT-induced pain.

### Activation of Gi DREADD reduces microglial IRF8 and IL-1β expression

Under neuropathic pain conditions, the transcription factor interferon regulatory factor 8 (IRF8) plays a significant role in microglial response (Masuda et al., 2012). Therefore, we tested the possibility that microglial Gi DREADD impacts neuropathic pain by modulating IRF8. To this end, we examined IRF8 expression after SNT with Gi DREADD activation (**Fig. 5A**). Our results showed that IRF8 is selectively expressed in CD11b^+^ microglia in the ipsilateral dorsal horn at POD3 after SNT (**Fig. 5B**). Pre-SNT activation of Gi DREADD by CNO significantly reduced SNT-induced IRF8 upregulation at POD3 (**Fig. 5B, C**). Western blot analysis revealed that SNT increased IRF8 expression for at least 5 days (**Fig. 5D**), and Gi DREADD activation reversed SNT-induced increase from POD1-3 days (**Fig. 5E**). IRF8 expression returned to comparable levels as control by POD5, which is also when mechanical allodynia became comparable to controls.

**Fig. 5.**
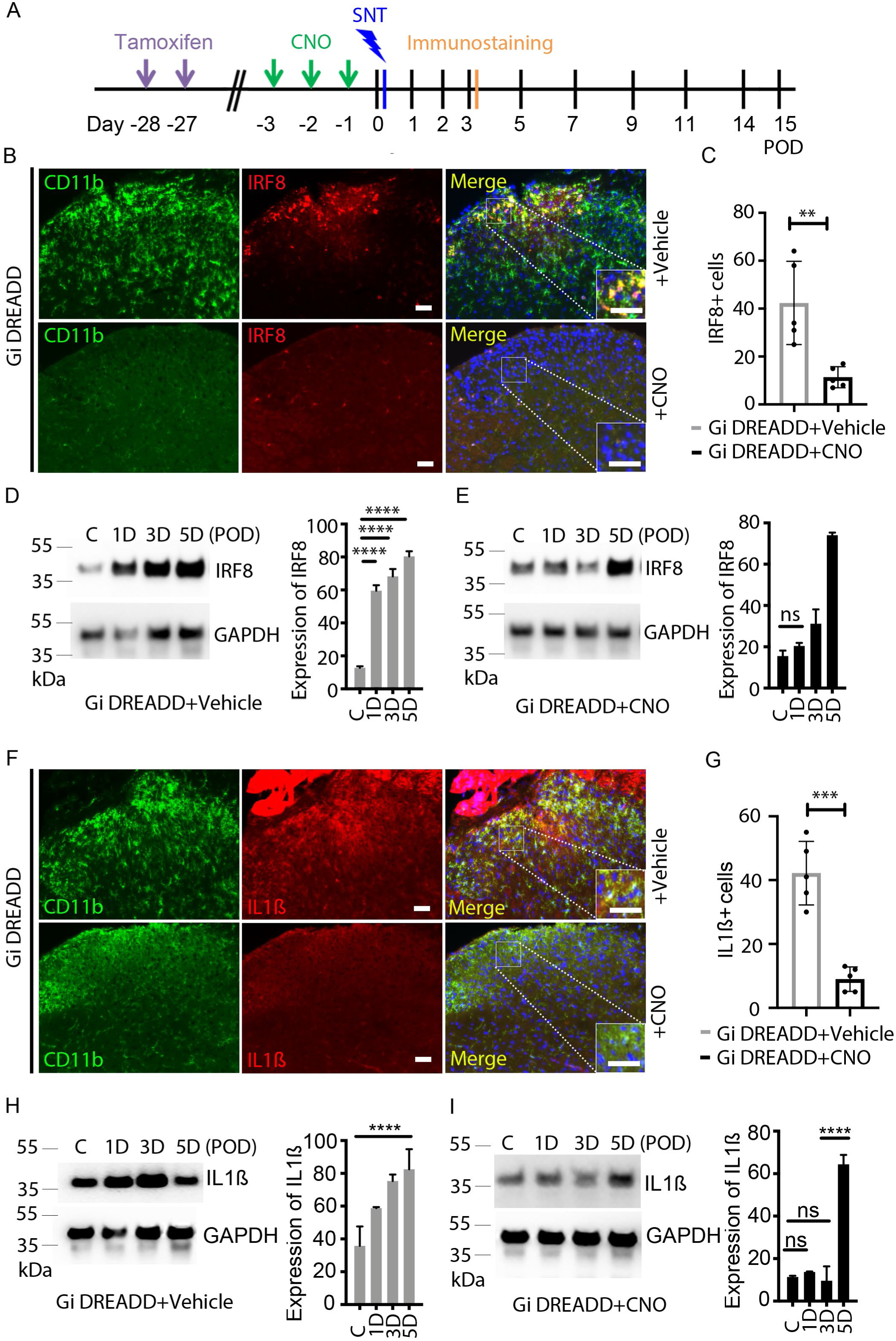
Microglial Gi DREADD activation inhibits IRF8 and IL1β expression. (A) Timeline of experimental procedures. (B) Immunofluorescence images showed that IRF8 (red) co-localized with CD11b ^+^ cells (green) and Dapi (blue) in the ipsilateral dorsal horn at POD3. The pre-SNT CNO treatment suppressed IRF8 expression. The insets show the magnified images of boxed area. Scale bar, 40 μm. (C) Pooled results show significantly reduced IRF8^+^ cells after CNO treatment as compared with that in vehicle group. n=5-8 mice/group, ** p < 0.01. (D) Representative Western blot images and quantification data showing that IRF8 expression in the L4-5 level of the dorsal horn increased after SNT n=5-8 mice/group, GAPDH was used as internal control. **** p < 0.0001. (E) Pre-SNT activation of microglial Gi DREADD by CNO suppressed the increase of IRF8 expression following SNT at POD1-3. n=5-8 mice/group. (F) Immunofluorescence images showing that IL1β (red) is co-localized with CD11b^+^ microglial cells (green) in the ipsilateral dorsal horn at POD3 following SNT. CNO treatment reduced SNT-induced IL1β expression in microglia. The insets show the magnified images of boxed area. Scale bar, 40 μm. (G) Pooled results show significantly reduced IL1β^+^ cells after CNO treatment compared with that in vehicle treated group. n=5-8 mice/group, *** p < 0.001. (H, I) Representative Western blot images and quantification data showing that IL1β expression in the L4-5 level of the dorsal horn was increased after SNT (H) which was suppressed by pre-SNT activation of microglial Gi DREADD by CNO (I). n=5-8 mice/group, GAPDH was used as internal control. **** p < 0.0001.

IRF8 is a critical transcription factor for the regulation of pro-inflammatory interleukin 1 beta (IL-1β), which is important for the pathogenesis of chronic pain (Ren and Torres, 2009; Gui et al., 2016). Thus, we investigated the possibility that the effect of Gi DREADD upon IRF8 also extended to IL-1β production by microglia. Consistently, we found that pre-SNT Gi DREADD activation significantly reduced IL-1β expression largely expressed in CD11b^+^ microglia (**Fig. 5F, G**). Moreover, Western blot analysis showed that up-regulation of IL-1β persists over POD1-5 after SNT (**Fig. 5H**), while CNO suppressed IL-1β upregulation through POD3 (**Fig. 5I**). Taken together, these results suggest that microglial Gi DREADD may exert its influence on chronic pain via the modulation of neuroinflammation, including IRF8 and IL-1β expression.

### Post-SNT Gi DREADD activation in microglia attenuates chronic pain and neuroinflammation

Our previous study using cell ablation approaches report that microglia are transiently required for the maintenance of neuropathic pain (Peng et al., 2016). Here, to directly assess the role of the post-SNT microglial Gi DREADD activation in the maintenance of neuropathic pain, we applied CNO at POD 3-5 after SNT in CX_3_CR1^creER/+^:R26^LSL-hM4Di/+^ mice (**Fig. 6A**). WT+CNO group and Gi DREADD+Vehicle group were examined as controls. Our result indicated that post-SNT activation of Gi DREADD in microglia reduced mechanical hypersensitivity in ipsilateral side, which lasted for 4 days in CX_3_CR1^creER/+^:R26^LSL-hM4Di/+^ mice but not in control groups (**Fig. 6B**). In addition, we found no sex difference in the inhibitory effect of post-SNT activation of Gi DREADD on chronic pain maintenance between male and female mice (**Fig. 6C**).

**Fig. 6.**
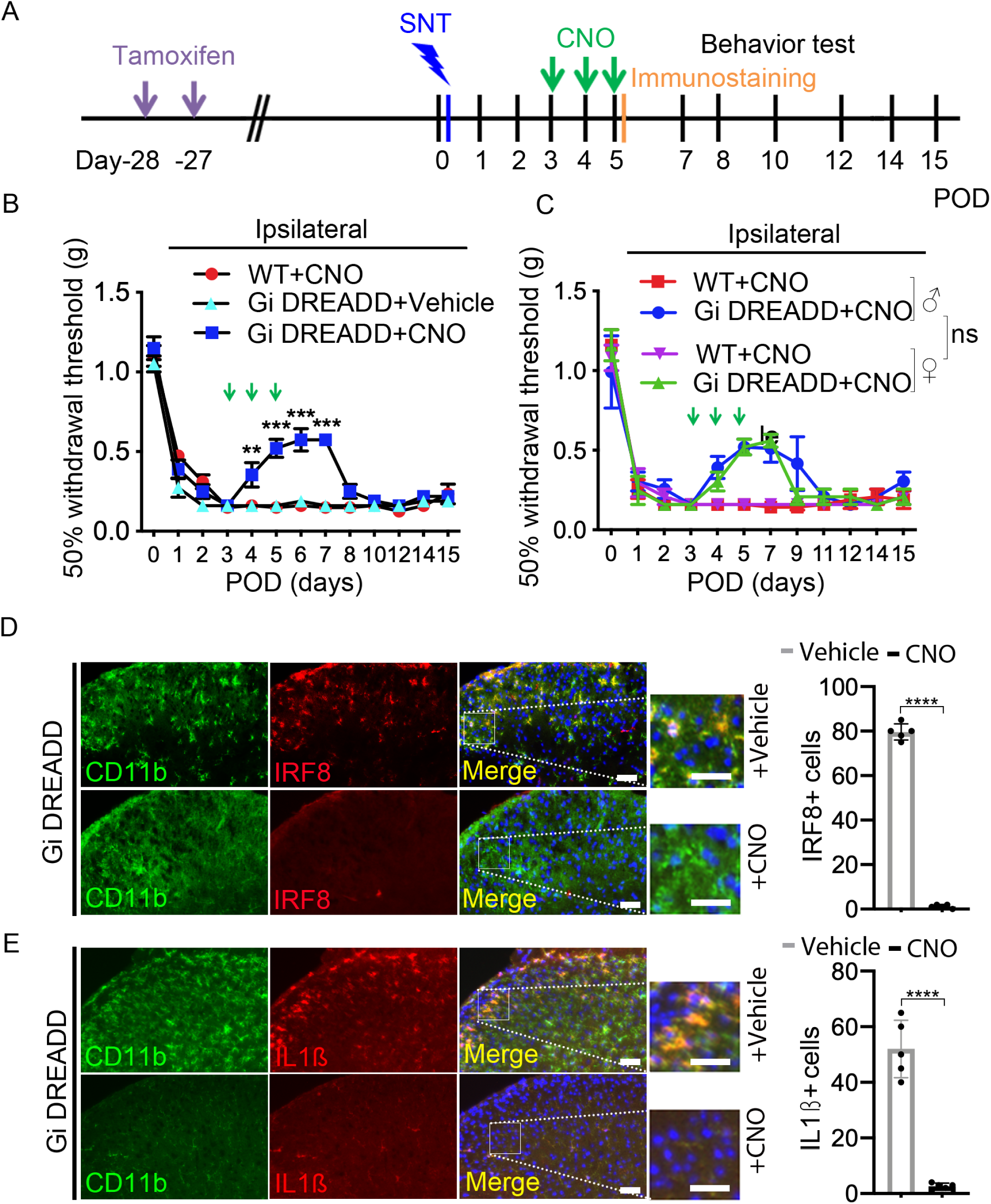
Post-SNT activation of microglial Gi DREADD reverses mechanical hypersensitivity and neuroinflammation induced by SNT. (A) Timeline of experimental procedures. (B) Behavioral test showing that post-SNT activation of microglial Gi DREADD attenuated mechanical hyperactivity in CX_3_CR1^creER/+^:R26^LSL-hM4Di/+^ mice (Gi DREADD+CNO) when compared to control groups (WT+CNO or Gi DREADD+Vehicle). Data represented as mean ± SEM, n=5 mice/group. ** p < 0.01, *** p < 0.001. (C) Post-SNT activation of microglial Gi DREADD attenuated mechanical hyperactivity in both male and female CX_3_CR1^creER/+^:R26^LSL-hM4Di/+^ mice (Gi DREADD+CNO). Data represented as mean ± SEM, n=5 mice/group. WT mice with CNO were considered to be controls. There was no difference between male and female groups. (D) Immunofluorescence images and summarized results show that IRF8 (red) co-localized with CD11b^+^ microglial cells (green) in the ipsilateral dorsal horn at POD5 following SNT. Post-SNT CNO treatment to activate microglial Gi DREADD reversed the increase of IRF8 expression by SNT. The insets show the magnified images of boxed area. Scale bar, 40 μm. n=5 mice/group, **** p < 0.0001. (E) Immunofluorescence images and summarized results show increased expression of IL1β which is co-localized with CD11b^+^ microglia cells in the ipsilateral dorsal horn at POD5 following SNT. CNO treatment reversed the increase of IL1β expression by SNT. The insets show the magnified images of boxed area. Scale bar, 40 μm. n=5 mice/group, **** p < 0.0001.

We also explored the effect of post SNT activation of microglial Gi DREADD on IRF8 and IL-1β expression using immunostaining. We found that microglial Gi DREADD activation at POD3-5 by CNO reduced the expression of IRF8 in CD11b^+^ microglia at POD5 after SNT compared to that of vehicle treatment (**Fig. 6D**). In addition, post-SNT activation of microglial Gi DREADD similarly reduced IL-1β expression in CD11b^+^ microglia (**Fig. 6E).** Thus, our results suggest that microglial Gi DREADD attenuates the maintenance of chronic pain likely through reduced neuroinflammation such as the downregulation of IRF8 and IL-1β in microglia.

### Gi DREADD reduces C-fiber-evoked field potential after SNT

Finally, we wanted to determine the effect of Gi DREADD upon C-fiber mediated nociceptive transmission which is intimately related to chronic pain induction and maintenance (Sandkuhler, 2007). To this end, we used *in vivo* recording in the spinal cord dorsal horn in anesthetized mice to study on C-fiber-evoked field potentials (Liu et al., 2017; Zhou et al., 2019). Three groups of mice were used: WT sham, WT SNT (POD3) and Gi DREADD SNT (POD3). For each experimental group, CNO was injected after 30 mins of baseline recording (**Fig. 7A**). We found that basal C-fiber-evoked field potentials were strongly decreased by CNO in Gi DREADD mice after SNT, but not in WT sham control or WT SNT group (**Fig. 7B, C**). After normalization to pre-drug values (baseline; -30-0 min before CNO administration), C-fiber-evoked field potential were maximally decreased by 44.8% ± 18.9% after CNO treatment (**Fig. 7C**). Interestingly, the inhibitory effect of Gi DREADD on C-fiber response in SNT group lasted less than one hour (**Fig. 7C**). These results indicate that nociceptive transmission after peripheral nerve injury is inhibited by the activation of microglial Gi DREADD.

**Fig. 7.**
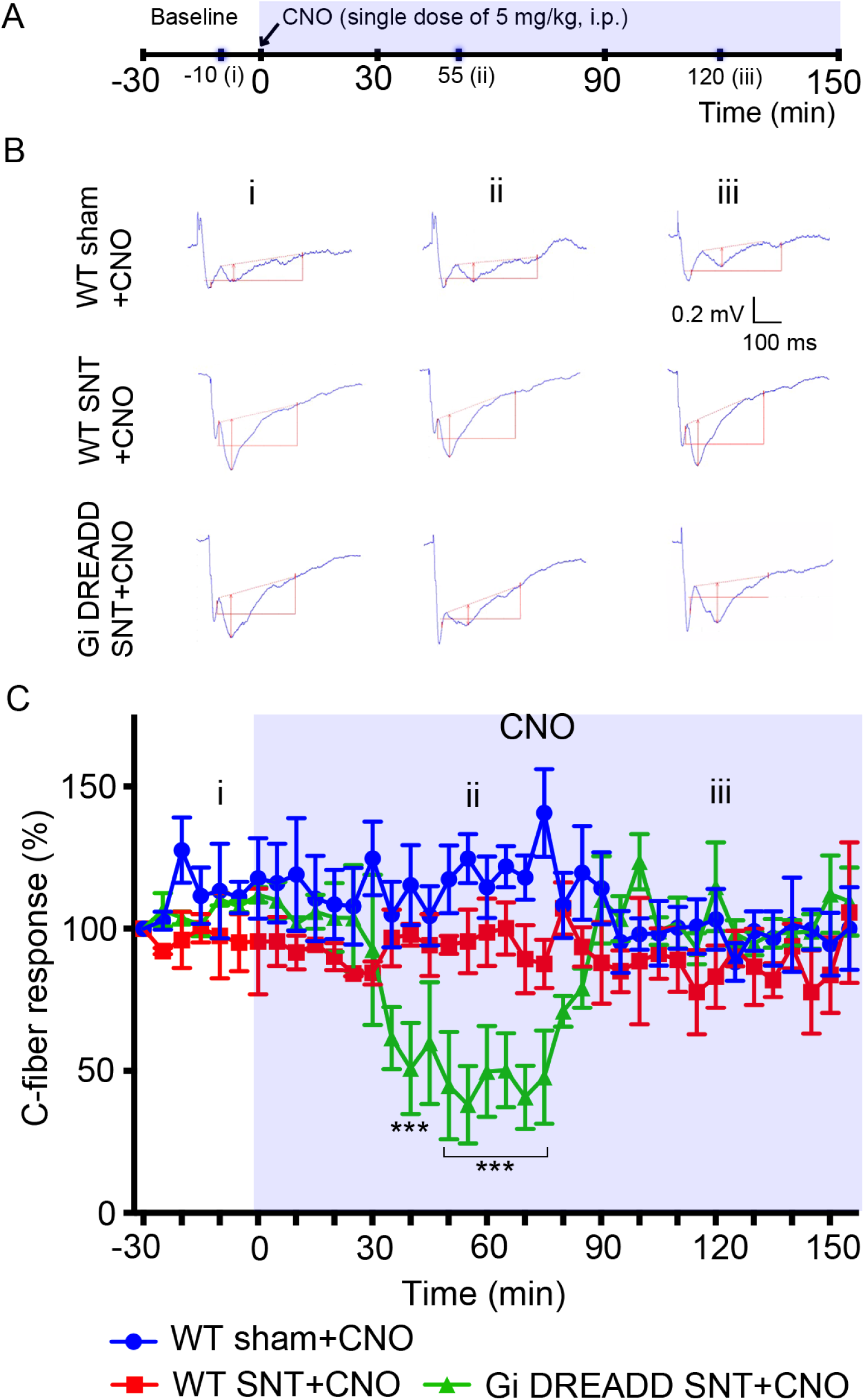
Microglial Gi DREADD activation attenuates C-fiber responses in the spinal dorsal horn after SNT. (A) Timeline of experimental procedures. (B) Representative traces of C-fiber field potentials from three groups (WT sham, WT SNT, and Gi DREADD SNT), recorded at baseline (i), after CNO treatment, 55 min (ii) and 120 min (iii). The amplitude of C-fiber-evoked response (red vertical line) was determined with parameter extraction software WinLTP. n=5 mice/group. Scale bars, 100 ms (X) and 0.2 mV (Y). (C) Pooled results showing the time course of C-fiber field potentials in response to the CNO application. CNO reduced C-fiber responses in Gi DREADD SNT group but not in control groups. C-fiber-evoked field potential was normalized to baseline. Data represented as mean ± SEM, n=5 mice/group. *** p < 0.001.

## Discussion

In this study, we used transgenic Gi DREADD approach as a novel tool to examine the role of microglia in neuropathic pain. Taking advantage of CX_3_CR1^creER/+^:R26^LSL-hM4Di/+^ transgenic mice, we were able to controllably activate Gi DREADD in microglia. Our results showed that microglial Gi DREADD attenuated SNT-induced microglial activation and chronic pain hypersensitivities. Mechanistically, we found that microglial Gi DREADD signaling reduced IRF8 upregulation, IL-1β production, and C-fiber evoked responses after peripheral injury. Together, these results demonstrate that microglial Gi signaling by chemogenetic manipulation attenuates chronic pain via inhibition of neuroinflammation.

Chemogenetic approaches like DREADD have only recently been employed to study glial function, in particular, astrocyte function in motor coordination and memory (Agulhon et al., 2013; Sciolino et al., 2016; Nam et al., 2019). Chemogenetic approaches have also been used to study oligodendrocyte control of neural activity (Ou et al., 2019). Finally, the Watkins’ group utilized DREADD approaches to study microglia in the spinal cord of rats via AAV9 mediated expression (Grace et al., 2016; Grace et al., 2018). By selectively expressing Gi DREADD under a CD68 promotor, the group elucidated microglial function in morphine-induced nociceptive sensitivity (Grace et al., 2016) and cytokine production in neuropathic pain (Grace et al., 2018). However, further study is needed to understand microglial DREADD downstream signaling in chronic pain behaviors. To this end, we employed a genetic mouse model to express Gi DREADD specifically in microglia. Our results are largely consistent with the study from Watkin’s group. However, our current study is exciting in several regards. First, this is the first genetic mouse model to manipulate microglia function *in vivo* using chemogenetic approaches. Second, our results suggest that expression of IRF8, IL-1β, and synaptic transmission can be regulated by Gi signaling during the development of neuropathic pain. Third, we showed that microglia are transiently required for the maintenance of neuropathic pain. Therefore, this study provides proof-of-principle that Gi signaling in microglia can suppress their pro-inflammatory activation and thus is a powerful tool to manipulate microglia function in chronic pain. One caveat is that Gi DREADD is also expressed in supraspinal microglia and thus may contribute to the analgesic effects of systemic CNO on SNT-induced pain. To circumvent this issue, local activation of microglial Gi DREADD is required in the future studies to delineate the specific function of spinal and supraspinal microglia in chronic pain.

In the present study we found that IL-1β, which is a critical mediator of neuropathic pain, was decreased by microglial Gi DREADD activation. These results demonstrate that pro-inflammatory cytokines produced by microglia can be regulated via Gi signaling. In addition, recent studies have concluded that IRF8 is a major regulator of IL-1β (Masuda et al., 2012). In the periphery, IRF8 plays a pivotal role in the immune system (Taniguchi et al., 2001; Honda and Taniguchi, 2006). We further demonstrated a link between microglial Gi DREADD signaling and IRF8 expression after SNT. Interestingly, ectopic expression of IRF8 causes marked upregulation of P2Y12 which is Gi-protein-coupled receptor in cultured microglia *in vitro* (Honda et al., 2001; Masuda et al., 2012). Therefore, this may provide the self-regulation mechanism for neuroinflammation in chronic pain. Collectively, our results suggest that expression of IRF8 and IL-1β are regulated by Gi signaling pathways which can modulate microglial activation and subsequent pain hypersensitivity. However, additional studies will be needed to further dissect how Gi signaling directly impacts IRF8 expression.

We also demonstrated with *in vivo* electrophysiology experiments that Gi DREADD significantly inhibited C-fiber-evoked field potentials following SNT. We observed the single dose of CNO reversed the nociceptive transmission from around 40 min post injection until at least 70 min post injection. These findings broadly match the known pharmacokinetic profile of CNO in the activation of DREADD (Jendryka et al., 2019). The mechanism is still unknown regarding how Gi DREADD activation acutely inhibits synaptic transmission. However, multiple lines of evidence have suggested that microglia can regulate neuronal activity. For example, microglia were able to dampen neuronal hyperactivity under seizure context (Eyo et al., 2014; Kato et al., 2016) or even under physiological conditions (Peng et al., 2019; Cserep et al., 2020). Particularly, selective microglial activation in the spinal cord was sufficient to facilitate synaptic strength between primary afferent C-fibers and spinal neurons (Clark et al., 2015; Zhou et al., 2019). Thus, considering the increased microglia reactivity under neuropathic pain condition, our results are plausible that Gi DREADD activation in microglia inhibited C-fiber synaptic transmission. Interestingly, we found that three doses of CNO were able to reduce chronic pain and the inhibitory effects lasted for several days. In addition, microglia Gi DREADD activation was accompanied by reduced neuroinflammation such as IRF8 and IL-1β. Thus, the longer term activation of Gi signal transduction via DREADD may lead to a biochemical cascade over multiple days. However, it still remains unknown how transient inhibition of synaptic transmission results in the long lasting effect on SNT-induced chronic pain after multiple times of microglial Gi DREADD activation.

Recent reports indicate that Gi DREADD can be successfully expressed and functional in non-human primates (Nagai et al., 2016). Therefore, it is plausible to explore the therapeutic potential of Gi DREADD in neuropathic pain. For example, AAV-CD68-hM4Di was used to express Gi DREADD in rat microglia (Grace et al., 2016). Using similar AAV vectors to express Gi DREADD in microglia in humans is possible, as they are commonly used for gene therapy in clinical patients (Naso et al., 2017). Moreover, consistent with its pharmacological inertness, CNO has been administered to humans without apparent toxic effect (Jann et al., 1994). However, CNO is not a drug that has been approved for use in humans by the Food and Drug Administration (FDA) (Lieb et al., 2019). Alternatively, FDA approved olanzapine is a second-generation atypical antipsychotic which is able to activate Gi DREADD (Weston et al., 2019). Nevertheless, this study is a proof-of-concept that Gi DREADD manipulation of microglia could influence neuropathic pain, suggesting its potential application for pain treatment in the long run.

## Acknowledgements

This work is supported by National Institutes of Health (R01NS088627, R21DE025689, R01NS112144) to L.J.W., and by a postdoctoral fellowship from the Mayo Clinic Center for Multiple Sclerosis and Autoimmune Neurology to T.C. We thank Dr. Anthony Umpierre and Dr. Aastha Dheer for critical reading of the paper and members of Wu lab at Mayo for insightful discussions.

## Author Contributions

M.H.Y and L.J.W. designed the studies. M.H.Y. performed the experiments and collected data. Y.U. L. assisted with some experimental design. K. L., T. C., J. Z. and M. X., assisted with analysis. L. Z assisted with *in vivo* electrophysiology recording. M.H.Y., D. B. and L.J.W. wrote and revised the manuscript.

